# Lexical parafoveal previewing predicts reading speed

**DOI:** 10.1101/2020.10.05.326314

**Authors:** Yali Pan, Steven Frisson, Ole Jensen

## Abstract

While reading is an essential human skill, many of the neuronal mechanisms supporting reading are not well understood. In spite of the reduced visual acuity, parafoveal information plays a critical role in natural reading^1^; however, it is strongly debated whether words are previewed parafoveally at the lexical level^2–9^. This is a key dispute for competing models on reading and of great significance since lexical retrieval is important for guiding eye movements. We found evidence for parafoveal previewing at the lexical level by combining a novel *rapid invisible frequency tagging* (RIFT)^10, 11^ approach with magnetoencephalography (MEG) and eye-tracking. In a silent reading task, target words of either low or high lexical frequency were tagged (flickered) subliminally at 60 Hz. The tagging response measured when fixating on the pre-target word reflected parafoveal previewing of the target word. We observed a stronger tagging response as early as 100 ms following fixations on pre-target words that were followed by target words of low compared with high lexical frequency. Moreover, the difference in tagging response with respect to target lexicality predicted individual reading speed. Our findings demonstrate that reading unfolds in the fovea and parafovea simultaneously to support fluent reading.

## Main

Humans have developed the remarkable skill of reading, allowing for efficient acquisition of information from busy pages or screens of text. Given the importance of written text for communication, individuals with reading disabilities are highly disadvantaged in modern society. Yet, we know little about the neuronal mechanism underlying natural reading. Our study aimed at answering if lexical information is retrieved for upcoming words in the parafovea (i.e., 2 to 5 degrees from the current fixation) during natural reading. We used a novel technique combining *rapid invisible frequency tagging* (RIFT) with magnetoencephalography (MEG) to provide neural insights on this question. In the present study, participants read 228 sentences in total (composed of two sets of sentences). All sentences were plausible and contained unpredictable target words of either low or high lexical frequency (see Supplementary information for sentence details). Word length for both pre-target and target words were matched with respect to low and high lexical frequency of the target words (Supplementary Table 1). Target words were flickered at 60 Hz throughout the reading of each sentence while the neuronal activity was measured by MEG (Fig. 1). When participants fixated on the pre-target word, the flickering target induced a reliable tagging response at 60 Hz, reflecting the neuronal resources associated with parafoveal previewing. Thus, this experimental design allowed us to investigate neural activity associated with lexical parafoveal previewing without interfering with natural reading.

**Figure 1.**
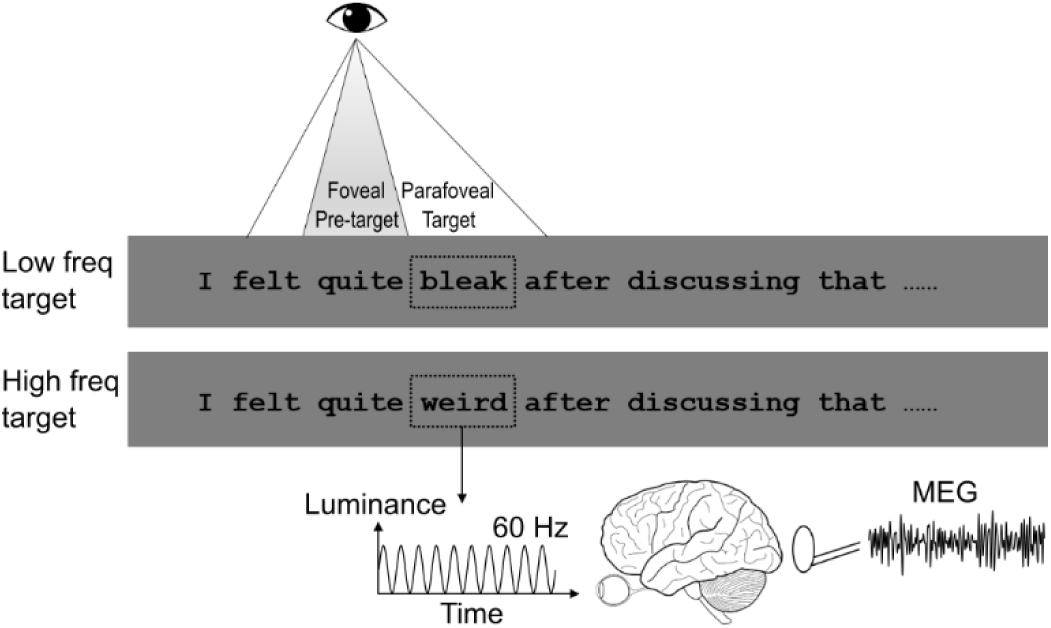
The reading task. Participants (n = 39) read sentences silently at their own pace, while eye-movements and brain activity were recorded. Each sentence was embedded with either a low or high lexical frequency target word (see dashed rectangle; not shown in the experiment). A Gaussian smoothed patch below the target word was flickered at 60 Hz continuously while the sentence was on the screen. This allowed us to measure the associated neuronal response using MEG: rapid invisible frequency tagging (RIFT). One-quarter of the sentences were followed by a simple yes-or-not comprehension question to ensure that participants read the sentences carefully. Freq., frequency.

In line with previous studies^2, 4, 5, 8, 9^, we did not find an effect of target word lexical frequency on pre-target first fixation durations (i.e. the duration of the *first* fixation on a word) (*t*_(38)_ = 0.17, *p* = 0.86, *d* = 0.03, two-tailed pairwise t-test, Fig. 2a). However, the target fixation durations were longer for low compared with high lexical frequency targets (*t*_(38)_ = 6.94, *p* = 3 × 10^−8^, *d* = 1.11, two-tailed pairwise t-test). This classic word frequency effect indicates the successful manipulation of target lexical frequency. We observed the same pattern using gaze durations (i.e. *total* duration of first-pass fixations on a word; Supplementary Fig. 1). These findings demonstrate that parafoveal lexical processing is not reflected in the eye movement data, at least in the well-controlled experimental studies.

**Figure 2.**
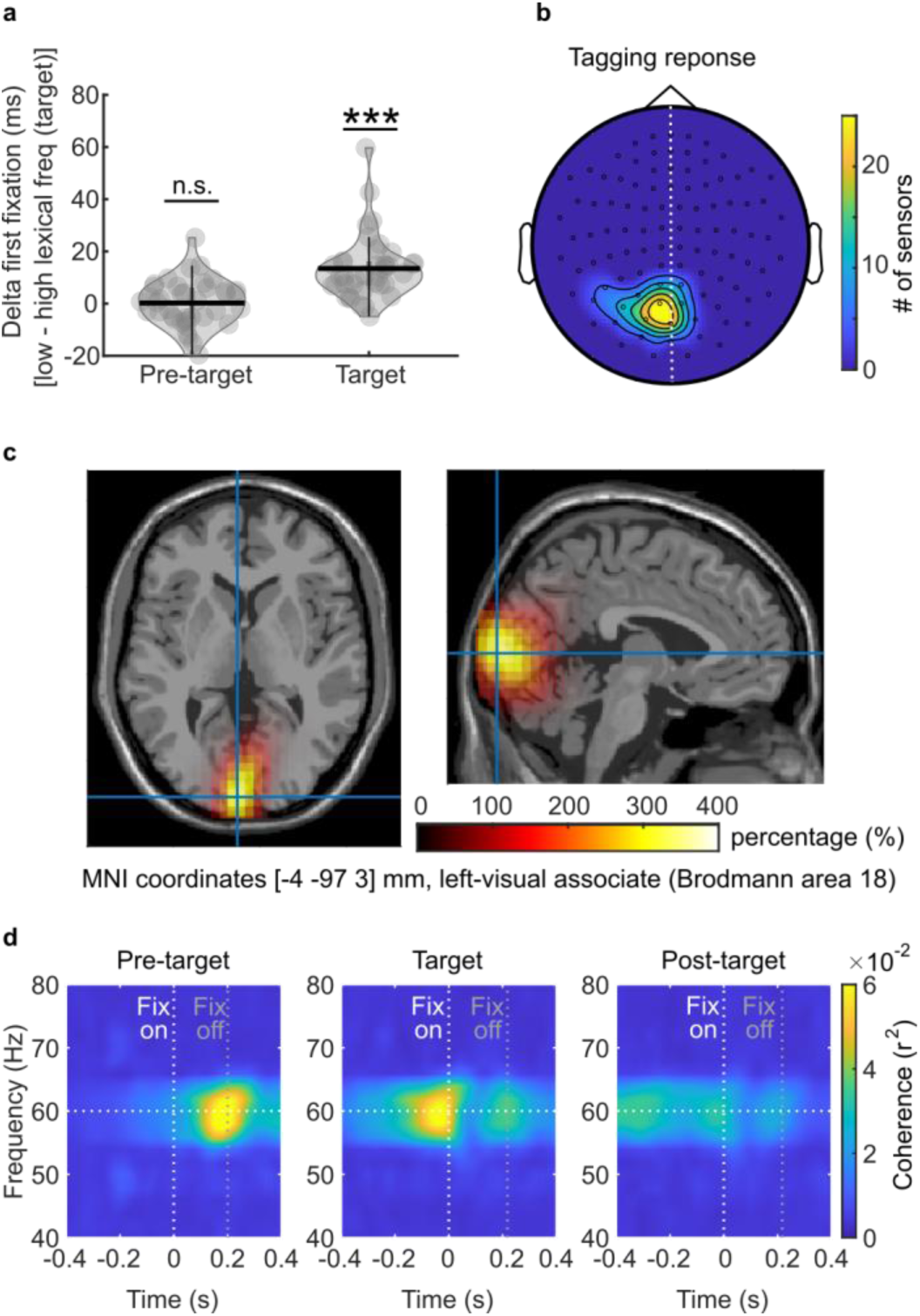
Eye-movement metrics and tagging responses. (**a**) The first fixation duration difference for pre-target and target words when comparing low-versus high lexical frequency target words. There was no difference in fixation times on pre-target words with respect to target lexicality (left). The fixation times were however longer for low than high lexical frequency target words (*** *p* < 0.001, n = 39, two-tailed paired t-test). The horizontal bar in violin plot indicates the mean value; each dot represents one participant. (**b**) Topography for sensors from all participants that showed stronger tagging responses during the pre-target period (flicker) compared with the baseline period (no-flicker). Twenty-six participants showed a robust RIFT response. The tagging response was quantified as the coherence between the tagging signal and the MEG signal. (**c**) Source modelling based on a beam-forming approach identified the neural sources of tagging response to the early visual cortex (Brodmann area 17, 18; relative change for pre-target 60Hz coherence compared with baseline). (**d**) The time-course of neuronal tagging response (coherence) during the pre-target (left panel), target (middle panel), and post-target fixation periods (right panel, n = 26). The analysis was time-locked to fixations onsets (white vertical lines; the grey vertical lines indicated the average fixation offsets). The strong coherence in the pre-target fixation interval captures brain activity associated with parafoveal target processing. Freq., frequency.

Next, we analyzed MEG data to uncover the brain activity associated with lexical processing before saccading to the target. In previous researches on spatial attention, we demonstrated that RIFT captured covert attention, reflected by stronger tagging responses for attended compared with unattended stimuli^10, 11^. Here we hypothesized that lexical parafoveal previewing would be observed as a modulation in the tagging response with respect to target lexical frequency during pre-target fixation. A measure of time-resolved coherence (r^2^) between the 60 Hz visual flicker and the brain activity was used to quantify the tagging responses (see Methods for details).

We found a strong tagging response over the left visual cortex sensors (Fig. 2b), reflecting the neuronal resources associated with parafoveal previewing. This was observed as a robust 60 Hz visual flicker-to-MEG coherence during pre-target fixations (caused by the target flickering in the parafovea) as compared with a baseline period (cross-fixation presented before sentence onset). This analysis was conducted pooled over target lexical frequency conditions. In 26 out of the 39 participants, one or more sensors had a significant tagging response (5.4 ± 4.0 sensors per participant, mean ± SD). The subsequent analyses were based on these participants and sensors. A source modelling approach revealed that the generators of this 60 Hz coherence difference were localized in the early visual cortex (Brodmann area 17, 18; Fig 2c). The time course of the 60 Hz coherence is shown in Fig. 2d. Note the robust 60 Hz coherence from the target word when fixating on the pre-target word. These results demonstrate that RIFT is a sensitive tool for measuring brain activity associated with parafoveal previewing during natural reading.

Next, we addressed if the RIFT was modulated by the lexical frequency of the target word. Our key finding was that the coherence at 60 Hz in the pre-target fixation was significantly stronger when followed by a low compared with a high lexical frequency target word (Fig. 3a; *t*_(25)_ = 2.20, *p* = 0.037, *d* = 0.43, two-tailed pairwise t-test). To ensure that the coherence in pre-target fixation was not contaminated by temporal smoothing from target fixation, the time window for averaging coherence was adjusted individually according to the shortest pre-target fixation duration (88.3 ± 8.9 ms across participants, mean ± SD). Furthermore, as the number of trials biases coherence magnitude, we subsampled the same number of trials for both conditions in each participant (by randomly selecting trials from the condition with more trials). In sum, we found evidence for a stronger neuronal response for low compared with high lexical frequency target words in the interval when participants fixated on the pre-target word. This finding supports lexical extraction in the parafovea.

**Figure 3.**
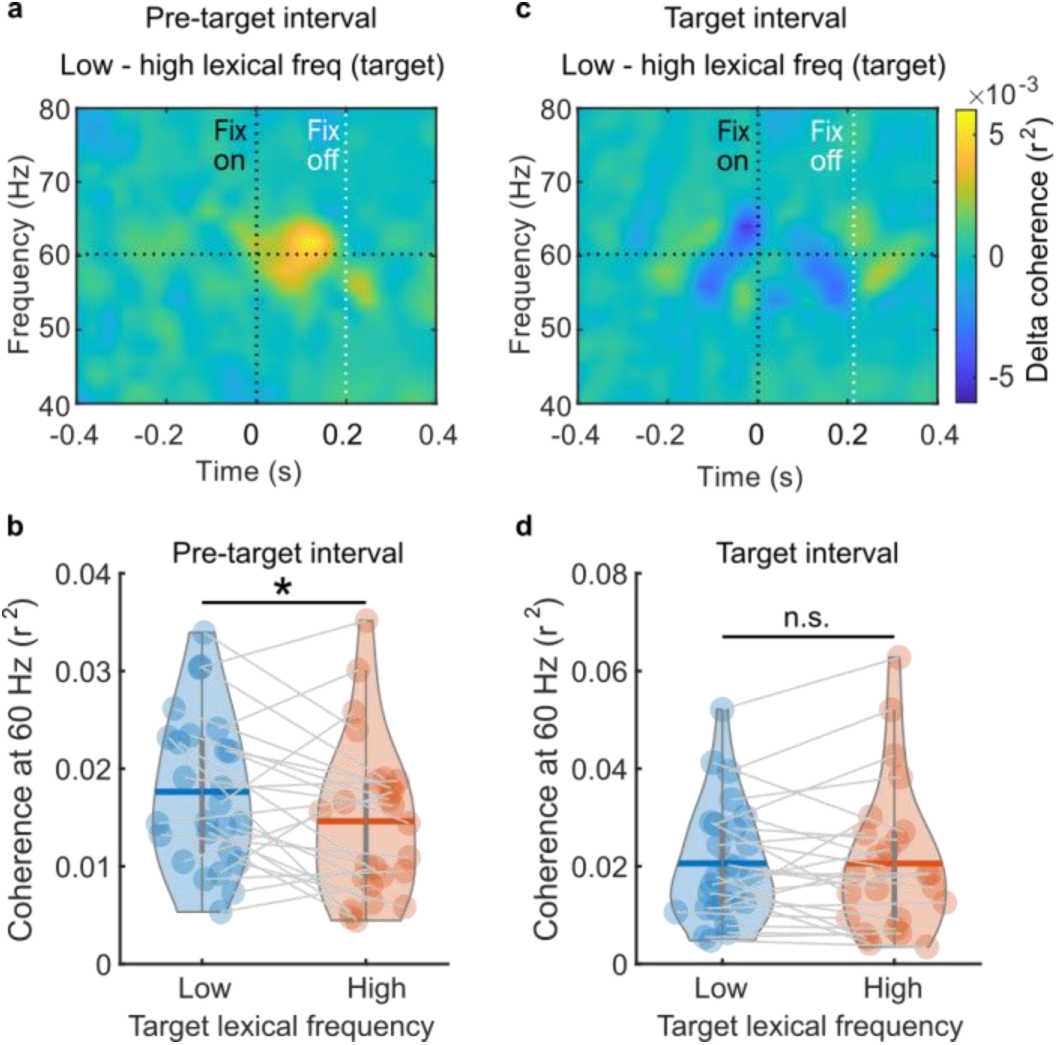
Neuronal activity associated with parafoveal processing. (**a**) Time-frequency representation of coherence differences during pre-target word fixations (low-minus high lexical frequency target words; time-locked to fixation onsets; n = 26). (**b**) The pre-target coherence at 60 Hz during pre-target word fixations for low-(blue) and high lexical frequency (orange) target words (n = 26, violin horizontal bar indicates mean value). The coherence was significantly stronger for low-compared with high lexical frequency target words (* *p* = 0.037; two-tailed paired t-test). (**c,d**) During target word fixations, the coherence was similar for low and high lexical frequency target words. Freq., frequency.

Did the lexical frequency of the target word affect onset latency for parafoveal previewing? The pre-target coherence onset latency was defined by the time to reach its half maximum (see Supplementary information). We then used a jackknife procedure to statistically evaluate if the coherence onset was significantly different for low compared with high lexical frequency target words. While no statistical difference was found for the combined sets of sentences (Supplementary Fig. 2a left panel, *t*_(25)_ = −1.69, *p* > 0.1, two-tailed pairwise t-test), we did however find a significant difference for the set with shorter pre-target and target words (2^nd^ sentences set; see Supplementary Fig. 2c left panel, *t*_(25)_ = −2.85, *p* < 0.01, two-tailed pairwise t-test). Importantly, this effect was already visible around the first 100ms of the pre-target fixation (72ms for low lexical frequency target; 116ms for high lexical frequency target). The null finding for the combined sentence set might be explained by the longer and more varying length of the pre-target and target words (Supplementary Fig. 2b). The faster onset for the low lexical frequency target indicates that parafoveal neuronal processing is modulated in time by upcoming lexical information as well, but this effect is only visible when pre-target and target words are relatively short.

Did lexical parafoveal previewing facilitate reading? To address this, we correlated the coherence difference with respect to lexical frequency of the target word when fixating the pre-target word with individual reading speed. Reading speed was quantified as the number of words read per second (calculated as the total number of words in a sentence divided by the respective sentence reading time). We found a positive correlation (Fig. 4; *r*_(25)_ = 0.45, *p* = 0.022, Spearman correlation), indicating that participants who captured more lexical information in the parafovea were also faster readers.

**Figure 4.**
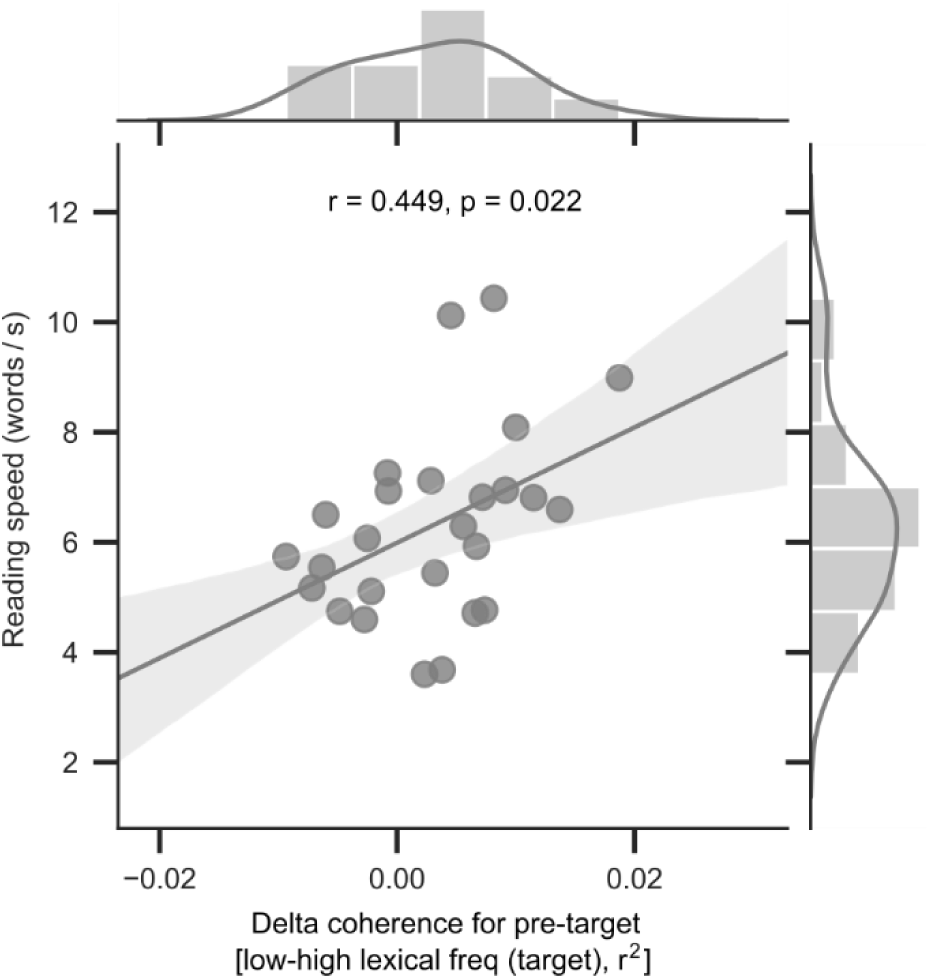
Relation between lexical parafoveal processing difference and individual reading speed. The coherence-difference with respect to target lexical frequency during fixations on pre-target words were derived per participant (see Fig. 3b). The reading speed was quantified as the number of words read per second. Each dot represents a participant. A Spearman correlation demonstrated a significant relation (n = 26). The shaded area indicates a 95% confidence interval. Histograms of individual reading speed and pre-target coherence differences are shown on the top and to the right. Freq., frequency.

In addition to the main analyses above, we also carried out two further analyses to rule out alternative explanations for our effects. First, we asked if the lexical parafoveal effect on the pre-target was coming from the target word but was smeared to the pre-target word (Fig. 3 c). Therefore, we tested whether the lexical frequency effect could also be observed during target fixations. As before, the averaging time window for the coherence was the minimum target fixation duration for each participant (87.1 ± 9.4 ms, mean ± SD). No significant coherence difference was found when fixating on low versus high lexical frequency target words (Fig. 3d; *t*_(25)_ = 0.01, *p* = 0.992, *d* = 0.002, two-tailed pairwise t-test). Presumably, the finding that the flickering did not affect target word viewing might be related to the fact that flicker sensitive photoreceptors (rod cells) mainly exist in the parafovea of the retina^12^. As a consequence, there would be reduced sensitivity to flickering when the target word is in foveal vision.

Next, we performed an initial landing position analysis to exclude the possibility that the lexical parafoveal effect was due to mislocated fixations or oculomotor error. If one fixation was intended for the target word, but the saccade undershot and landed on the pre-target word, then the lexical frequency effect of the target word should have been observed for the eye movements to the pre-target word. This was not the case in our data as no target lexical frequency effect was found for pre-target fixation durations (Fig. 2a). Furthermore, we showed that saccade landings on the pre-target words were highly accurate (−1.1 ± 1.9 letters with respect to the centre of the pre-target word, mean ± SD) and were similarly distributed as saccades to all words (Supplementary Fig. 3). The lexical frequency of the target word also did not affect the pre-target landing position (*t*_(25)_ = 0.35, *p* = 0.727, *d* = 0.07, two-tailed pairwise t-test). In addition, the coherence during the pre-target word fixations did not correlate with the landing position on the pre-target word (*r*_(25)_ = 0.16, *p* = 0.420; Spearman correlation).

Our findings shed new light on the long-standing debate between models arguing for either serial attention shift^13, 14^ or parallel graded processing ^15, 16^. Both models regard spatial attention as important for reading but they differ on whether more than one word can be accessed simultaneously at the lexical level. Parallel graded models predict that the lexical frequency of both foveal and parafoveal words can be processed simultaneously. This idea is challenged by the fact that only a few correlation-based corpus studies found lexical parafoveal effects in eye movement data^6, 7^, while more well-controlled experimental studies did not^2, 4, 5^ (and neither did our eye movement results, Fig. 2a). But these conclusions are based on eye movement studies, we found neural support for parallel graded models: lexical parafoveal effects were observed within 100ms from pre-target onset (Fig. 3b). This finding does not seem to be compatible even with the most temporally compressed version of a serial model, in which the attentional shift and a significant amount of lexical parafoveal previewing occur during saccadic programming^17^. According to such an account, the neuronal computations that need to be completed before parafoveal previewing can even begin - retina-brain lag, visual encoding, foveal lexical processing, and shift of attention -take far too long (around 150ms) to also accommodate the extraction of lexical information from the next parafoveal word. Moreover, evoked response studies based on EEG^18^ and MEG^19^ provide evidence for lexical frequency effects for foveally fixated words no earlier than 100 ms. Our results showed a lexical frequency effect for the parafoveal word at around 100 ms, which supports the idea of parallel processing of foveal and parafoveal information. In particular, we provide evidence that parafoveal lexical frequency modulates neural excitability in support of a parallel model of reading.

One might ask why lexical parafoveal previewing is reflected in the neuronal responses (Fig. 3b) but not in fixation durations (Fig. 2a)? This discrepancy indicates that two systems are at play simultaneously: covert and overt attention. The fixation duration on a foveal word (pre-target) captures overt attention, while the RIFT, which reflects the steady-state response to the parafoveal word (target), captures covert attention. However, these two parallel processes might not get equal attentional resources. Because the parafoveal word attracts less attention than the foveal word, its activation level might be insufficient to affect the current foveal saccade planning and fixation duration. Other studies using a rapid simultaneous visual presentation paradigm also show that words can be identified in parallel without rising to a high level of awareness^20^. Nevertheless, even though readers might not be consciously aware of their parafoveal previewing, it does accelerate individual reading as shown in our correlation analysis (Fig. 4). Taken together, our study shows that natural reading involves the simultaneous processing of several words, providing novel neural evidence for the idea that “readers are parallel processors”^21^.

The neuronal response reflecting previewing was observed in the early visual cortex. This might be a surprise, as functional Magnetic Resonance Imaging studies have localized lexical frequency to e.g. the visual word form area^22^. According to interactive processing theories, higher-level lexical information interacts with lower-level visual information during word recognition^18, 23^. Thus, lexical frequency information extracted in the parafovea could direct visual attention covertly. Increased spatial attention will boost tagging responses^10, 11^, resulting in stronger coherence for the pre-target word followed by a low compared with a high lexical frequency target word.

Our results show that RIFT is a new powerful technique to investigate parafoveal reading. A classic paradigm in this field is the gaze-contingent boundary task developed by Keith Rayner in 1975^24^. In this task, parafoveal information is manipulated by changing the target word while saccading to it^4, 25^. This approach allows for manipulating parafoveal previewing and has made great contributions to studies on parafoveal previewing^26^. However, the approach is limited, as changing the target word inevitably interferes with natural reading. Fixation related potentials based on EEG is another method used in reading paradigms^27^. While this approach has provided important insights by demonstrating a lexical frequency effect for foveal word recognition on the N1 component^18, 28^, it fails to provide conclusive results with regard to parafoveal lexical processing ^4, 5^. The novel approach we present in this study based on subliminal frequency tagging allows for capturing brain activity associated with parafoveal previewing during natural reading.

It will be important for future studies to use the RIFT to investigate other factors of parallel information capture during reading. Can parafoveal information be extracted at the semantic level? Another direction is investigating the primary determinants of reading proficiency in relation to parafoveal previewing. For instance, previewing at the phonological level identified using gaze-contingent paradigms has been found to reflect reading proficiency^29^. We found that the neuronal signature of lexical previewing predicted reading speed, which could be used as a potential indicator to diagnose reading disorders such as dyslexia. Besides, some researchers argue that dyslexia is due to spatial processing problems in the magnocellular visual pathway^30^. Our frequency tagging approach could be helpful to understand the underlying neural mechanism of dyslexia in relation to the allocation of spatial attention and various factors of previewing including parafoveal previewing at the phonological level. In sum, the present study demonstrates that RIFT is a powerful tool for investigating natural reading, and provides novel neural evidence for lexical parafoveal previewing in support of parallel graded models of reading.

## Methods

### Participants

Our study recruited forty-three participants (28 females), aged 22 ± 2.6 (mean ± SD), right-handed, with normal or corrected-to-normal vision, and without a neurological history or language disorder diagnosis. Four of them were excluded from analysis due to poor eye tracking or falling asleep during the recordings, which left thirty-nine participants (25 females). The University of Birmingham Ethics Committee approved the study. The participants provided written informed consent and received £15 per hour or course credits as compensation for their participation.

### Reading task

Each trial started with a central fixation cross on a grey screen centre presented for 1.2 − 1.6 s. Then followed by a square (1 degree wide) presented 2 degrees to the right of the middle of the screen left edge. A gaze of at least 0.2 s on this square triggered sentence onset. The square was replaced by the first word of the sentence. The text was presented in the equal spaced Courier New font, and each letter occupied 0.35 visual degrees (Fig. 1). After reading the sentence, participants were instructed to fixate on a square below the screen centre for 0.1 s to trigger the sentence offset. One-quarter of the trials were followed by a simple yes-or-no comprehension question to ensure careful reading. Two sets of sentences, consisting of respectively 142 and 86 sentences, were divided into 7 blocks. Each block took approximately 7 minutes to read and was followed by a rest for at least 1 minute (see Supplementary Information for sentence sets details).

### Eye-tracking

Each session began with a nine-point calibration and validation test. After every three trials, we performed a one-point drift checking test. If a participant failed to pass drift checking or was unable to trigger sentence onset through gazing, the nine-point calibration and validation test was conducted again. The eye-tracking error was limited to below 1 visual degree both horizontally and vertically.

### Visual projection and frequency tagging

All visual stimuli were presented by a PROPixx DLP LED projector (VPixx Technologies Inc., Canada) at a 1440 Hz refresh rate that provided a stable 60 Hz tagging signal (for a detailed description about how the projector works, please see Supplementary information). A Gaussian smoothed patch underlying the target word was flickered at 60 Hz during the whole sentence presentation. A custom made photodiode (Aalto NeuroImaging Centre, Finland) was attached to the right-below corner of the screen to record tagging signal from a square whose luminance was kept the same as the flickering patch.

### Coherence calculation

To measure the tagging response associated with target word processing, coherence was estimated between the MEG sensors and the tagging response of the photodiode (for MEG sensor selection see Supplementary information). First, 1s segments were filtered using a phase preserving, two-pass, Butterworth bandpass filters (4^th^ order) with a hamming taper. The centre filter frequencies were from 40 to 80 Hz in steps of 2 Hz with a 10 Hz frequency smoothing. For each frequency step, the analytic signals were determined by the Hilbert transform which then was used as the input for coherence at time point ***t***:

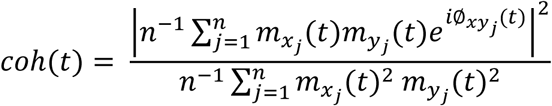

Where *j* is the trial, *n* is the number of trials, *m_x_(t*) and *m_y_(t)* are the time-varying magnitude of the analytic signals from respectively a MEG sensor and a photodiode, *Ø_xy_(t)* is the phase difference as a function of time between them. A time-frequency coherence representation was obtained as applied in Fig. 2d and Fig. 3a and 3c (for details of the source modelling of the MEG data see Supplementary information).

### Data and code availability

The data in this study, as well as the codes to generate the associated figures, will be available upon request from the corresponding author.

## Acknowledgements

We thank Dr. Federica Degno and Prof. Simon Liversedge for sharing the second sentence set, Dr. Geoffrey Brookshire and Yang Cao for feedbacks on the manuscript. This work was supported by the following grants to O.J.: the James S. McDonnell Foundation Understanding Human Cognition Collaborative Award (grant number 220020448), Wellcome Trust Investigator Award in Science (grant number 207550), and the BBSRC grant (BB/R018723/1) as well as the Royal Society Wolfson Research Merit Award.

## Author Contributions

Y.P. and O.J. devised and designed the experiments with assistance from S.F., Y.P. made the sentences with assistance from S.F, Y.P. programmed and conducted the experiments, Y.P. carried out the analyses with assistance from O.J. and S.F., Y.P., O.J. and S.F. wrote the paper together.

---

**Supplementary Table 1.** Characteristics of words in the experimental materials (1^st^ set).

| Measure | Pre-target | Target |  |  |  | Post-target |
| --- | --- | --- | --- | --- | --- | --- |
|  | Length | Low frequency | High frequency | Length | Position | Length |
| Mean | 6.1 | 5.3 | 95.3 | 5.8 | 6.7 | 6.7 |
| SD | 1.5 | 4.5 | 135.5 | 0.8 | 2.3 | 1.7 |

Low (<10) and high lexical frequency (>30) target words are reported in terms of the total CELEX frequency per million<sup>1</sup>. Length is the number of letters in a word. Position refers to the location in a sentence where the target word is presented, measured in words. The sentences were $11.2 \pm 2.1$ words long. For details about words in the 2<sup>nd</sup> sentence set, please see Degno et al., 2019<sup>2</sup>.

**Supplementary Figure 1.**
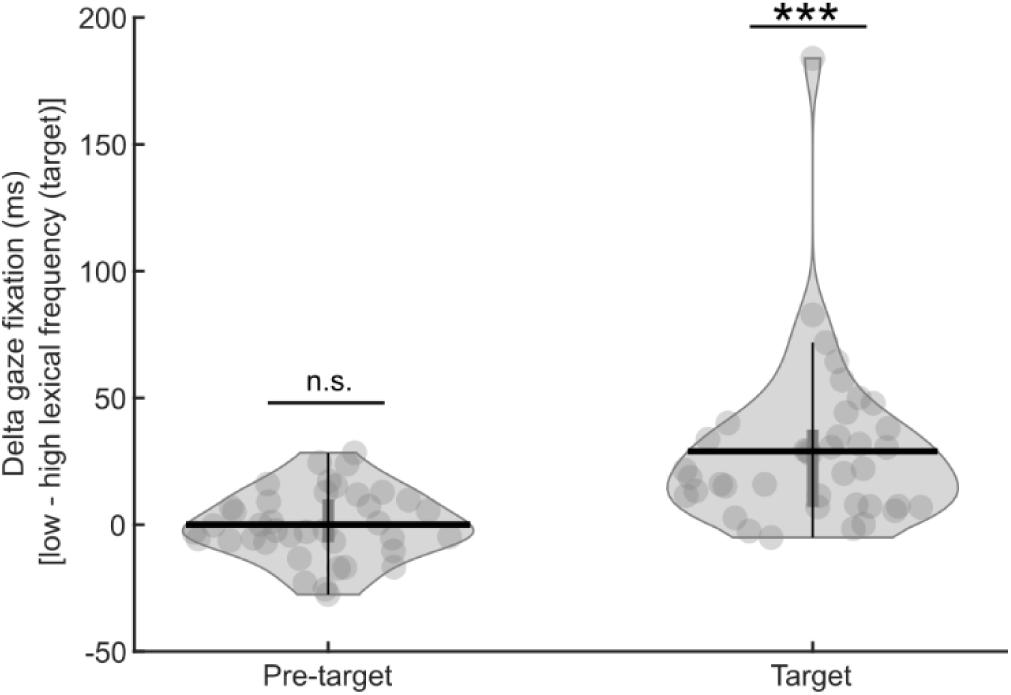
Gaze-duration difference for pre-target and target words. Gaze duration refers to the sum of all first-pass fixations for a given word. The differences for low-minus high-lexical frequency of the target words were calculated. A significant lexical frequency effect was only observed for target words, with longer gaze durations for low-compared with high-lexical frequency (*** *p* < 0.001, n = 39, two-tailed paired t-test). We observed no significant gaze duration difference for pre-target words *(p* = 0.977, n = 39, two-tailed paired t-test). The horizontal bar in the violin plot indicates mean value; each dot represents one participant. This finding supports the conclusion of Fig. 2a.

**Supplementary Figure 2.**
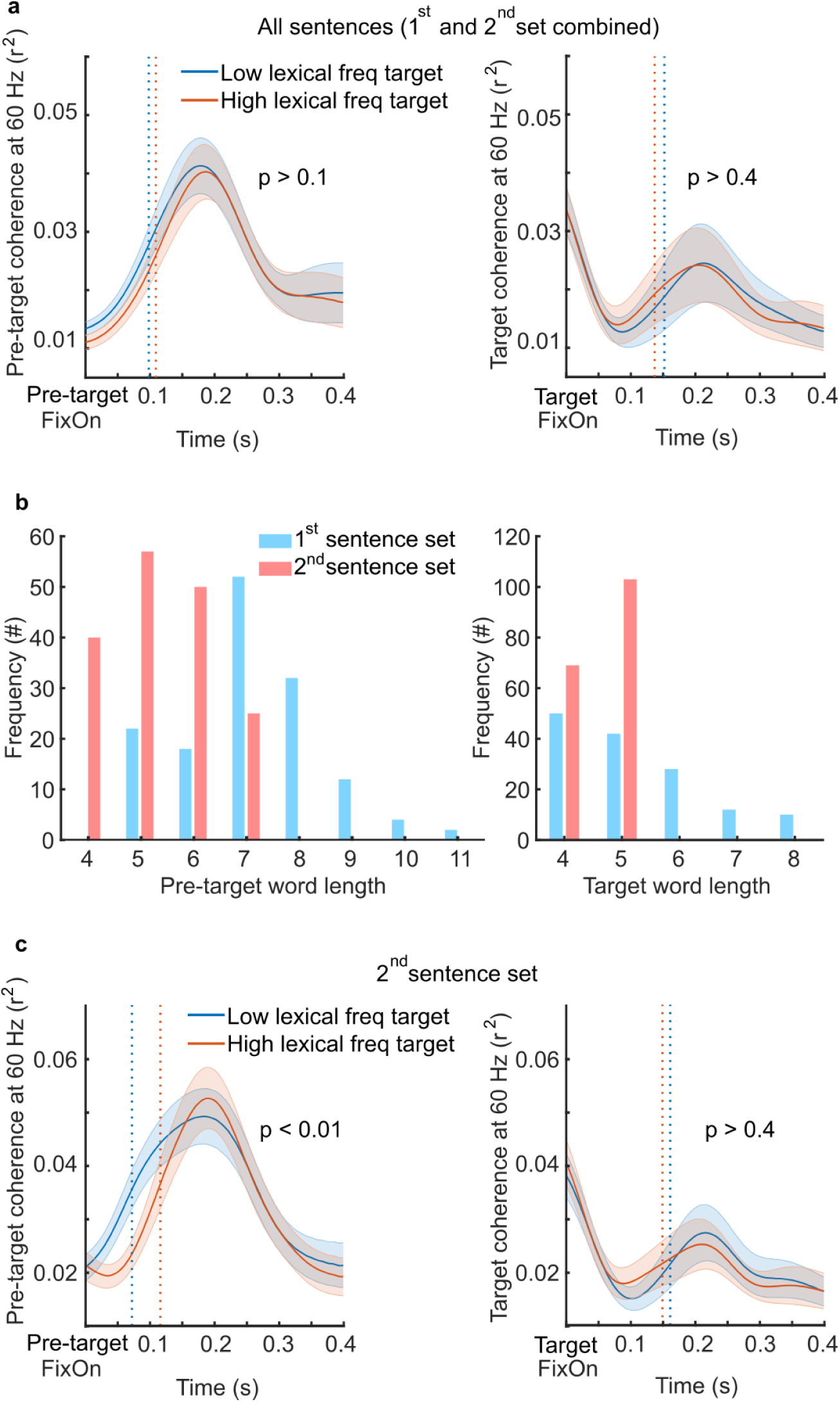
Pre-target coherence onset latency was modulated by parafoveal target lexical frequency in the short-words sentence set (2^nd^ Set). A jackknife-based method was used to identify the onset latency for 60 Hz coherence on the group level. Here onset latency refers to the time when coherence reached half maximum, denoted by the dotted lines. (**a**) Each participant read two sets of sentences; for the combined sentence set, no significant difference was found in either pre-target fixation (left panel, *p* > 0.1, n = 26, two-tailed paired t-test) or target fixation (right panel, *p* > 0.4, n = 26, two-tailed paired t-test). (**b**) The histogram of word length showed that in the 2^nd^ sentence set (in red), both pre-target and target words were shorter compared with the 1^st^ sentence set (in blue). (**c**) For the 2^nd^ sentence set, a significant coherence onset difference was found in pre-target fixation (left panel, *p* < 0.01, n = 26, two-tailed paired t-test), but not in target fixation (right panel, *p* > 0.4, n = 26, two-tailed paired t-test). (All analyses in the study were based on the combined sentence set except this one).

**Supplementary Figure 3.**
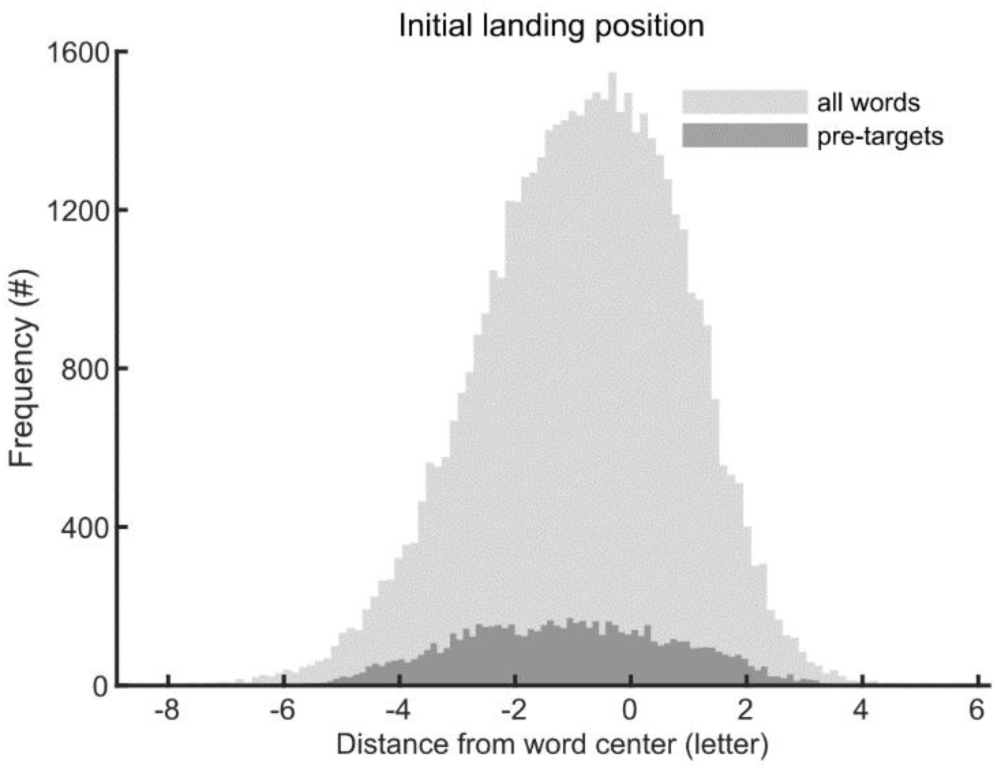
Initial landing position for pre-target words and all words. Landing positions for the first fixation on words were estimated in the unit of ‘letter’ aligned according to the word centre. The distribution in dark grey is the landing position error for pre-target words (−1.1 ± 1.9, mean ± SD); the landing position error for all words (−0.8 ± 1.8, mean ± SD) can be found in light grey. This demonstrates that the landing position of fixations was around ∼1 letter to the left of the centre of the fixated words.

## Supplementary Information

### Sentence material

#### 1^st^ sentence set

We constructed 142 sentences embedded with 71 target word pairs (low/high lexical frequency). For each sentence, the pre-target, target, and post-target words were in the same structure as adjective + noun + verb. For each target pair, two different sentence frames were made, and each participant read both target words embedded in these two different frames. For example, for the target pair **waltz/music** (low/high lexical frequency), one participant read version A, another one read version B (see below, targets are in bold for illustration, but not in the real experiment).

~~~
A. Mike thought this difficult **waltz** received lots of criticism.
It was obvious that the beautiful **music** captured her attention.
B. Mike thought this difficult **music** received lots of criticism.
It was obvious that the beautiful **waltz** captured her attention.
~~~

The sentences in version B were made from version A by circular shifting the first and second half of the sentences. For both versions, no more than 3 successive sentences were from the same target lexical frequency condition.

#### 2^nd^ sentence set

This sentence set was adapted from Degno et al., 2019^2^. We removed sentences that contained the same pre-target or target words as in the 1^st^ sentence set, which left 86 sentences. Each sentence was embedded with two target words from the same lexical frequency condition (see below, version A contained two low lexical frequency targets, while version B contained two high lexical frequency targets).

~~~
A. I felt quite **bleak** after discussing that really **risky** subject with Paul.
B. I felt quite **weird** after discussing that really **nasty** subject with Paul.
~~~

Each participant read either version A or B. The same control for sentence presentation was counterbalanced as in the 1^st^ set.

### Pre-tests for target predictability and sentence plausibility

We carried out behavioural pre-tests for the 1^st^ sentence set only; for pre-test results for the 2^nd^ set please see Degno et al., 2019^2^. The participants in the pre-tests did not participate in the MEG session.

#### Predictability

A cloze test was performed using the sentence fragment up to but not including the target word. Participants were asked to read the sentence silently and then write down the first word/s that came to mind to complete the sentence. Example:

~~~
Mike thought this difficult
~~~

If a target word (e.g. *waltz* or *music)* was generated by less than 10% of the participants, then the target was considered unpredictable. Twenty-two participants (1 male) took part in this pre-test, and 6 target words turned out to be highly predictable. These 6 target words were replaced and the predictability test was conducted again with another 22 participants (3 males). None of the target words were judged predictable.

#### Plausibility

Participants were instructed to rate how plausible (or acceptable) each sentence was. Plausibility was rated on a 7 point scale (see examples below). Sentences in the experiment were supposed to be highly plausible, in order to occupy the full range of the scale, we constructed 142 filler sentences with low plausibility, half of which were of middle plausibility (e.g., sentence 1 below) and another half were implausibility (e.g., sentence 3 below). In this example, sentence 2 was used in the experiment.

|  | Implausible |  |  |  |  | Plausible |  |
| --- | --- | --- | --- | --- | --- | --- | --- |
|  | 1 | 2 | 3 | 4 | 5 | 6 | 7 |
| 1. Kate said that she saw lots of stars twinkling in the sky at noon. | 1 | 2 | 3 | 4 | 5 | 6 | 7 |
| 2. It was obvious that the beautiful music captured her attention. | 1 | 2 | 3 | 4 | 5 | 6 | 7 |
| 3. Sarah used an adhesive to glue the traffic and just made it to the wedding. | 1 | 2 | 3 | 4 | 5 | 6 | 7 |
| ..... | 1 | 2 | 3 | 4 | 5 | 6 | 7 |

For version A (n = 22, 2 males, 2 invalid for incomplete and careless responses), the plausibility rating for the experimental sentences 5.7 ± 0.5 (mean ± SD), which was significantly higher than the filler sentences that were of low plausibility (two-tailed pair-wised *t*_(19)_ = 23.98, *p* < 0.001). For version B (n = 23, 4 males), the rating was 5.5 ± 0.8 (mean ± SD), which was also significantly higher than the filler sentences (two-tailed pair-wised *t*_(22)_ = 16.68, *p* < 0.001). These results showed that all the sentences in both versions were highly plausible.

### Procedure

Participants were seated comfortably in the MEG gantry, 145 cm away from the projection screen in a dimly-lit magnetically shielded room. One-line sentences were presented on a middle-grey screen using Psychophysics Toolbox -3^3^. Every sentence started at the same position: two degrees to the right of the middle of the screen left edge and was presented on the vertical centre. Words were displayed in black font colour with an equal-spaced Courier New font (size 22). Each letter and space between two words occupied 0.35 visual degrees. In total, no sentence was longer than 27 visual degrees horizontally. Participants were instructed to read each sentence silently at their own pace and to keep their head and body as stable as possible during the MEG session. Eye movements were acquired during the whole session. One-quarter of the sentences were followed by a simple yes-or-no comprehension question to ensure a careful reading. Participants put their left and right index fingers of the right hand on a two buttons response box to respond to the questions. The mapping between yes-or-no responses and button presses was counterbalanced over participants. All participants answered the questions with high accuracy (95.4% ± 4.7%, mean ± SD).

### Rapid Invisible frequency tagging (RIFT)

#### Flickering of the target word

To flicker the target word, we added a rectangular patch underneath the target. The width of the patch was the width of the target word plus the spaces on both sides. The height of the patch was 1.5 times the word height. The target word was placed in the centre of this rectangular patch. All pixels within the patch were flickered at 60 Hz by multiplying the luminance of the pixels with a 60 Hz sinusoid (the modulation depth was 100%). Typically, the patch was perceived as indistinguishable from the middle-grey screen background, which made it invisible to participants. To reduce the visibility of the patch edges during saccades, a Gaussian smoothed transparent mask was applied on top of the flickering patch. The mask was created by a two-dimensional Gaussian function:

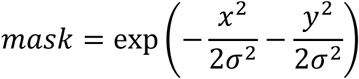

where, *x* and *y* are the mesh grid coordinates for the flickering patch, and σ is the x and y spread of the blob with σ = 0.02 degrees. By applying a Gaussian smoothed mask, the flickering patch was hardly perceived. Only three out of all the thirty-nine participants noticed the flickering patch according to a questionnaire after the MEG session.

#### Projector

To generate the rapid invisible frequency tagging, sentence stimuli were presented with a refresh rate up to 1440 Hz using a PROPixx DLP LED projector (VPixx Technologies Inc., Canada). This was done by presenting the sentence stimuli repeatedly in four quadrants on the stimulus computer screen (1920 × 1200 pixels resolution) with a refresh rate of 120 Hz. For each quadrant, the stimuli were coded in RGB three colour channels. The projector interpreted these 12 colour channels (3 channels × 4 quadrants) as 12 individual grayscale frames and projected them onto the projector screen separately in rapid succession. Hence, the refresh rate for stimuli presentation was 1440 Hz (120 Hz x 12).

### Data acquisition

#### MEG

MEG data were acquired using a 306-sensor TRIUX Elekta Neuromag system with 204 orthogonal planar gradiometers and 102 magnetometers (Elekta, Finland). The data were band-pass filtered online using anti-aliasing filters from 0.1 to 330 Hz and then sampled at 1,000 Hz. We used a Polhemus Fastrack electromagnetic digitizer system (Polhemus Inc, USA) to digitize the locations for three bony fiducial points: the nasion, left and right preauricular points. Then, four head-position indicator coils (HPI coils) were digitized: two coils were attached on the left and right mastoid bone and another two were on the forehead with at least 3 cm distance in between. Furthermore, at least 200 extra points on the scalp were acquired for each participant in order to spatially co-register the MEG source analysis with individual structural MRI image. After preparations, participants were seated upright under the MEG gantry with the back rest at a 60°angle.

#### Eye movements

The eye-tracker (EyeLink 1000 Plus, SR Research Ltd, Canada) was placed on a wooden table in front of the bottom edge of the projector screen. The distance between the eye-tracker camera and the centre of the participant’s eyes was 90 cm. It was used throughout the whole experiment to acquire horizontal and vertical positions of the left eye as well as the pupil size. Eye movements were sampled at 1,000 Hz. We also placed one pair of electrodes above and below the right eye (vertical electrooculogram, EOG) and another pair to the left and right of the eyes (horizontal EOG) to provide additional measures for ocular and eye-blink artefacts.

#### MRI

After MEG data acquisition, we acquired the T1-weighted structural MRI image using a 3-Tesla Siemens PRISMA scanner (TR = 2000 ms, TE = 2.01 ms, TI = 880 ms, flip angle = 8 degrees, FOV = 256×256×208 mm, 1 mm isotropic voxel). Out of all the thirty-nine participants, three dropped out of the MRI acquisition, one of them showed robust tagging responses at the sensor level. For this participant, the MNI template brain (Montreal, Quebec, Canada) was used instead in later source analysis.

### MEG data analyses

The data analyses were performed in MATLAB R2019b (Mathworks Inc, USA) by using the FieldTrip^4^ toolbox (version 20200220) and custom-made scripts.

#### Pre-processing

The MEG data were band-pass filtered from 0.5 to 100 Hz using phase preserving two-pass Butterworth filters. First, the MEG segments were extracted from -0.5 to 0.5 s intervals aligned with the first fixation onset for pre-target, target, and post-target words, respectively. Only segments with fixation durations ranging from 0.08 to 1 s entered further analyses. Segments with too short or too long fixations were discarded. We also extracted 1 s long baseline segments aligned with the presentation onset for the cross-fixation, which was the period before sentence onset. Next, the MEG data were demeaned by removing the linear trend and the mean value. After removing malfunctioning sensors (0 to 2 sensors per participant), these segments entered an independent component analysis^5^ (ICA). Before the ICA, a PCA approach was applied to reduce the rank of the data to 30 components. Next, the components related to eye blinks, eye movements, and heartbeat were rejected. Finally, we manually inspected all these segments to further identify and remove any segments that were contaminated by excessive noise like ocular, muscle, or movement artefacts.

#### RIFT response sensor selection

To identify the MEG sensors that showed reliable tagging responses, we compared the 60 Hz coherence during pre-target segments with the coherence during baseline segments. We used a non-parametric statistics method named Monte-Carlo to estimate the significance for the coherence difference. This method was developed by Maris et al., 2007^6^, and implemented in the Fieldtrip^4^ toolbox. Both pre-target and baseline segments were 1 s long and were aligned with the first fixation onset for pre-target words and the onset for baseline cross-fixation separately. The pre-target segments were constructed by pooling the target lexical frequency conditions together. Several previous RIFT studies from our lab observed robust tagging responses from the visual cortex for visual flickering stimuli ^7–10^. Therefore, only MEG sensors in the visual cortex (52 planar sensors) entered this sensor selection procedure.

Here, we regarded pre-target and baseline segments as two conditions in the coherence calculation. For a given MEG sensor and photodiode combination, coherence at 60 Hz was estimated over trials for each condition. Therefore, one coherence value was obtained for each condition. Then, we calculated the z-statistic value for this coherence difference between pre-target and baseline using the following equation (for details please see Maris et al., 2007^6^):

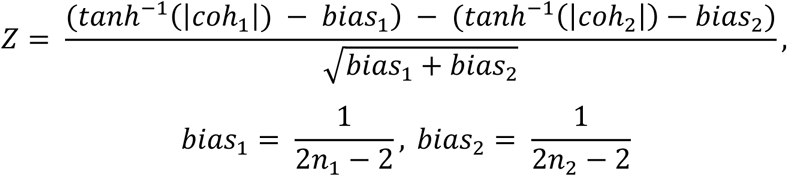

where *coh_1_* and *coh_2_* denote the coherence value for pre-target and baseline condition, *bias_1_* and *bias_2_* is the term used to correct for the bias from trial numbers of pre-target (*n*_1_) and baseline condition (*n*_2_). So, all trials from the pre-target and baseline conditions were used. After obtaining the z statistic value for the empirical coherence difference, a permutation procedure was conducted to estimate the significance probability.

We randomly shuffled the trial labels between pre-target and baseline conditions for 10,000 times. During each permutation, coherence values for both conditions were re-computed so that a z statistic value was estimated for the coherence difference. After all the shuffles, a null distribution for z-values was established. If the empirical z-value was larger than 99% of the null distribution, which meant that the coherence difference between pre-target and baseline was larger than zero at the 0.01 significance level, this sensor was considered to have robust tagging responses. This sensor selection procedure was performed for every sensor in the visual cortex (52 planar sensors in total). Twenty-six out of all the thirty-nine participants showed robust tagging responses at one or more sensors (5.4 ± 4.0 sensors per participant, mean ± SD; Fig. 2b). For each participant, the coherence values were averaged over all tagging response sensors to obtain an averaged coherence.

#### Source analysis for RIFT

In order to localize the neural sources that were coherent with the photodiode signals during RIFT, a beamforming approach was performed using Dynamic Imaging Coherent Sources (DICS) ^11^ implemented in the FieldTrip^4^ toolbox. The DICS technique enabled us to calculate the source estimates in the frequency domain with a focus on 60 Hz, which was the RIFT frequency. The beamformer was based on adaptive spatial filters derived for each grid in the discretized brain volume. In this source analysis, only participants with robust tagging responses were included (n = 26; see Fig. 2b).

A semi-realistic head models was constructed using a procedure developed by Nolte in 2003^12^, which uses spherical harmonic functions to fit the brain surface. We first aligned the individual structural MRI image with the MEG data. This was done by spatially co-registering the three fiducial anatomical markers from the head shape digitization during the MEG session (nasion, left and right ear canal). For one participant whose MRI image was unavailable, the MNI template brain was used instead. Next, this aligned MRI image was segmented into a grid. Then, we prepared the single-shell head model based on the segmented MRI image

The individual source model was constructed by inverse-warping a 5 mm spaced regular grid in the MNI template space to each participant’s segmented MRI image in the native space. This regular grid was from the Fieldtrip template folder and was constructed before doing the source analysis. In this way, the beamformer spatial filter was constructed on the direct grid that mapped to the MNI template space.

The Cross-Spectral Density (CSD) matrix was calculated between all the MEG sensor combinations and between the MEG sensor and the photodiode combination at 60 Hz in the whole 1 s time window, with a smoothing of 4 Hz using the hanning taper. For each participant, we estimated CSD matrices for both the pre-target and baseline segments.

Next, a common spatial filter was computed based on the individual single-shell head model, source model, and CSD matrices using DICS. This spatial filter was applied to both the pre-target and baseline CSD matrices for coherence computation. This was done by normalizing the magnitude of the summed CSD between the MEG sensor and the photodiode by their respective power. After the grand average over participants, the relative change for pre-target coherence was estimated as the ratio between coherence difference and baseline coherence ((*coh_pretarget_ -coh_baseline_)/coh_baseline_*). Finally, this source analysis localized the RIFT neural sources to the left-visual associate, Brodmann area 18, MNI coordinates [-4 -97 3] (see Fig. 2c, n = 26).

#### Jackknife-based method for onset latency

We used a leave-one-out jackknife-based method^13^ to assess the onset latency difference for the pre-target and target coherence separately with respect to the target lexical frequency (Supplementary Fig. 2). During each iteration, one randomly chosen participant was left out, and the averaged coherences for low and high target lexical frequency condition were calculated over the remaining participants. Then, the coherence onset latencies were computed for both conditions. Here, the onset latency was defined as the time point when the averaged coherence value reached its half-maximum *(coh_min_ + (coh_max_ - coh_min_)/2*). We computed the onset latency difference by subtracting the onset latency for the low target lexical frequency condition from the high condition. After twenty-six iteration (n = 26), onset latency differences from all these subsamples were pooled together to estimate a standard error *(SD*) using the following equation:

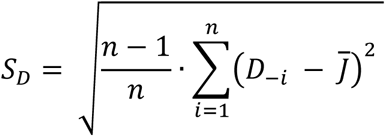

where the 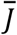 is the averaged onset latency difference over all the subsamples, *D_-i_* is the coherence difference obtained from the subsample when participant *i* was left out, *n* is the number of participants. We also computed the onset latency difference from the overall sample set (without leaving any participant out), and divided it by the *S*_*D*_ to obtain its *t*-value. A standard t table provided the statistical significance for the coherence onset latency difference between high and low target lexical frequency conditions. This procedure was done for both pre-target and target segments to conduct a statistical test for the coherence onset latency difference between both target lexical frequency conditions as shown in Supplementary figure 2a and 2c. (for details about the jackknife-based method for measuring onset latency differences, please see Miller et al., 1998^13^).

### Statistical information

All the t-tests in this study were two-sided pairwise student’s t-tests and were conducted in R^14^.

## Notes

### Competing Interest Statement

The authors have declared no competing interest.

